# The Mouse Epididymal Amyloid Matrix: A Mammalian Counterpart of a Bacterial Biofilm

**DOI:** 10.1101/2023.11.15.567275

**Authors:** Caitlyn Myers, Georgia Rae Atkins, Johanna Villarreal, R. Bryan Sutton, Gail A. Cornwall

**Author notes:** Department of Microbiology and Immunology, Institute of Biomedicine, University of Gothenburg, Gothenburg, Sweden (Caitlyn Myers). Department of Obstetrics, Gynecology and Reproductive Sciences, University of Pittsburgh School of Medicine, Pittsburgh, PA.

## Abstract

The mouse epididymis is a long tubule connecting the testis to the vas deferens. Its primary functions are to mature spermatozoa into motile and fertile cells and to protect them from pathogens that ascend the male tract. We previously demonstrated that a functional extracellular amyloid matrix surrounds spermatozoa in the epididymal lumen and has host defense functions; properties not unlike that of an extracellular biofilm that surrounds and protects a bacterial community. Here we show the epididymal amyloid matrix also structurally resembles a biofilm by containing eDNA, eRNA, and mucin-like polysaccharides. Further these structural components exhibit comparable behaviors and perform functions like their counterparts in bacterial biofilms. Our studies suggest that nature has used the ancient building blocks of bacterial biofilms to form an analogous structure that nurtures and protects the mammalian male germline.

## Introduction

The epididymis is a highly convoluted tubule connecting the testis to the vas deferens. Its primary functions are to mature spermatozoa into motile and fertile cells and to protect them from pathogens that ascend the male tract. Because spermatogenesis occurs after self-tolerance is established, spermatozoa are naturally antigenic (1,2). Therefore, the epididymal lumen is an immunologically unique compartment. It maintains a tolerant environment for spermatozoa to enable their maturation into functional cells while at the same time can mount a host defense response against pathogens. Precisely how the epididymis carries out and coordinates these two critical functions remains unclear.

We previously showed that the mouse epididymis contains a functional extracellular amyloid matrix that is distributed throughout the lumen and is closely associated with the maturing spermatozoa(3). The epididymal amyloid matrix includes cystatin C and several CRES (cystatin-related epididymal spermatogenic) subgroup family members including CRES, CRES2, CRES3, and cystatin E2(4). Unlike cystatin C, CRES subgroup members lack consensus sites for cysteine protease inhibition suggesting unique biological roles(5). Indeed, CRES, CRES2, CRES3, and cystatin E2 are highly aggregation-prone, each forming amyloids with distinct assembly properties *in vitro* (4). Further, CRES3 and CRES can cross-seed suggesting interactions between subgroup members, possibly by the formation of heterooligomers, may regulate epididymal amyloid matrix assembly (6). We recently demonstrated that CRES amyloids and the epididymal amyloid matrix have host defense functions and adopt different amyloid structures (matrix, film, fibrils) with different antimicrobial functions (bacterial trapping, killing, promotion of ghost-like bacteria) depending on the pathogenicity of the bacterial strain (7). Thus, the shape-shifting properties of the amyloid matrix allow for a plasticity that is integral for its functions.

The contribution of several amyloidogenic CRES subgroup members to the assembly of a mammalian host defense structure is strikingly like the amyloidogenic curli family of proteins that organize the assembly of *E. coli* biofilms (8). Curli member B (CsgB) serves as a nucleator to initiate curli A (CsgA) amyloid assembly at the bacterial cell surface forming the infrastructure of the extracellular biofilm (9). A primary function of biofilms is to nurture the bacteria within them by providing a safe environment which is stable and rich in nutrients(10). Biofilms also protect bacteria from antibiotics, pathogens, and other harmful factors making them especially efficient host defense structures(11). Based on these structural and functional similarities, we hypothesized the epididymal amyloid matrix may be a mammalian counterpart of a bacterial biofilm. However, rather than nurturing and protecting a bacterial colony as a biofilm would do, the epididymal amyloid matrix nurtures/matures and protects the male germline.

The structural components of bacterial biofilms have long been studied as potential therapeutic targets for dispersing bacteria during infections (12). Biofilms are composed of bacteria and extracellular polymeric substances (EPS) that can include amyloid, extracellular DNA (eDNA), extracellular RNA (eRNA), and polysaccharides (13). The many roles these components perform in the formation and function of biofilms are still an area of intense study. Amyloids serve an essential function as a structural scaffold interacting with other extracellular components in the biofilm (14). They also can contribute to the adherence of the biofilm to substrates and perform a protective role by trapping phage particles (15,16). eDNA can have several roles in biofilms, including in its formation by templating amyloid assembly (17), regulation of its maturation(18), maintenance of overall structure(19–21), cell adhesion (22), horizontal gene transfer (23), and protection from antibiotics(24). More recently, eRNA has also been shown to be a component of some biofilms, and like eDNA, contributes to overall biofilm structure including its distinctive viscoelastic properties (25). Further, several noncoding RNAs have been shown to regulate biofilm assembly by mediating curli production and exopolysaccharide synthesis (26). Polysaccharides, often mucopolysaccharides/glycosaminoglycans, are the major component of the EPS providing the “glue” important for the structure and adhesive properties of the biofilm. These long linear chains of sugars also stratify the bacteria within the community (27,28) and form a physical barrier or protective coat that can promote a tolerogenic immune response for bacteria (27,29–31). Their ability to protect eDNA/eRNA from degradation by nucleases suggests a close association between polysaccharides and nucleic acids within the biofilm (32). In addition, polysaccharides including sulfated forms, like eDNA, have been shown to template amyloid assembly (33,34).

Here, we carried out studies to characterize the structural components of the mouse epididymal amyloid matrix to determine if it has the properties of a bacterial biofilm and contains extracellular DNA, RNA and polysaccharides. Further, we determined if these structures perform similar functions and exhibit comparable behaviors as their counterparts in bacterial biofilms. Establishing the component parts of the epididymal amyloid matrix is key for elucidating the mechanisms of its assembly, host defense functions, and interactions with its resident cells, the spermatozoa. This knowledge is essential for developing much needed therapies for male infertility including that as a result of luminal occlusions formed following bacterial infections (35). Discovering parallels between the epididymal amyloid matrix and biofilm structures/functions also could prove insightful for identifying basic mechanisms essential for cell/species survival, including those mediated by environmentally driven epigenetic modifications. We demonstrate that eRNA, eDNA, and mucin-like polysaccharides are key components of the epididymal amyloid matrix and play critical roles in maintaining its overall structure. Together, our studies reveal how nature has preserved the ancient building blocks of biofilms to construct a structure that nurtures/protects the mammalian male germline.

## RESULTS

### eDNA and eRNA are components of the epididymal amyloid matrix

We used differential centrifugation to isolate fractions containing distinct populations of the epididymal amyloid matrix (pellets 2 and 4 (P2, P4)) from the mouse caput and corpus-cauda (from here on referred to as cauda) epididymis (Fig 1A, B). Our previous studies suggest the amyloid matrix is assembling in the most proximal part of the epididymis (caput) as sperm enter from the testis, forming a highly branched matrix that transitions into fibrillar arrays in the distal/cauda epididymis (7). The P2 and P4 fractions may represent precursor (early condensates) and more mature amyloid assemblies (matrix, fibrils), respectively, within each region. The P2 and P4 amyloid-containing fractions were spread on a slide and stained with Sytox Green, Hoechst, and TOTO-3, dyes commonly used to visualize nucleic acids, and thioflavin S (ThS), a fluorescent dye that recognizes the cross-β sheet structure of amyloid. As shown in Fig 1C, D both P2 and P4 amyloids from the caput epididymis and P4 amyloid from the cauda epididymis were stained with all three nucleic acid dyes and ThS indicating the presence of nucleic acids in the amyloid matrix. Although the nucleic acid dyes are routinely used as indicators of DNA, they can also bind RNA. Pretreatment of the caput P2 and P4 amyloids with 5% SDS, which helped expose the amyloid as indicated by increased ThS fluorescence, did not affect staining suggesting nucleic acids are part of an amyloid infrastructure that is resistant to, or protected from, SDS. Similar results were observed in the cauda epididymis treated with 70% formic acid. Formic acid instead of SDS was needed to facilitate exposure of the amyloid consistent with it being a more mature structure in this epididymal region. We next performed experiments in which the caput and cauda amyloid samples were costained with ThS and propidium idodide (a nucleic acid stain). The merged fluorescent images show that nucleic acid colocalizes with amyloid suggesting it is part of the amyloid infrastructure (Fig 2).

**Figure 1.**
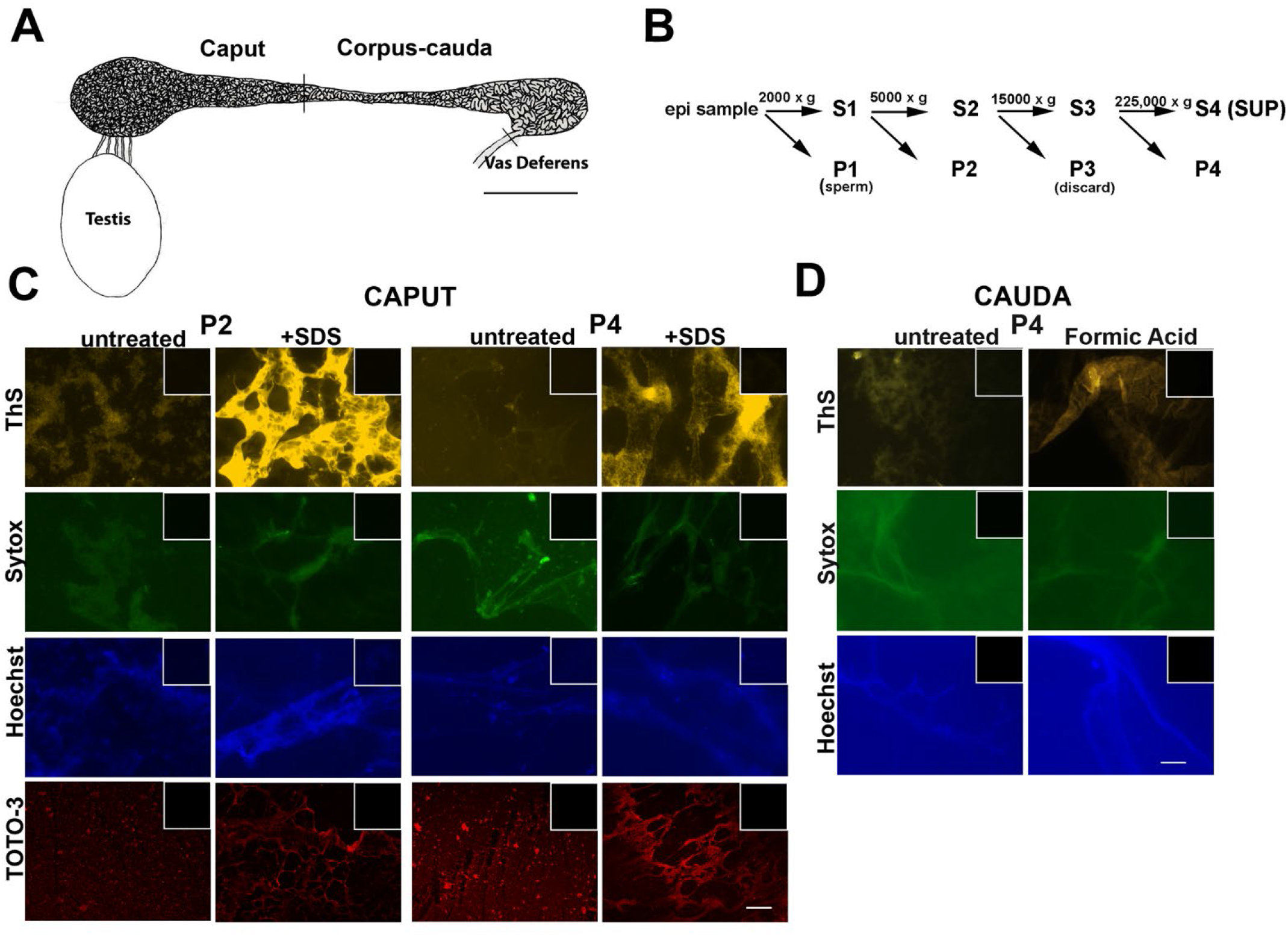
Extracellular nucleic acids are present in the epididymal amyloid matrix. **A)** Schematic diagram of the rodent epididymis showing the caput and corpus-cauda (cauda) regions and its connections to the testis and vas deferens. Scale bar, 3 mm. **B)** Differential centrifugation protocol used to isolate fractions enriched in amyloids of different molecular mass from the caput and cauda epididymal lumen. S, supernatant; P, pellet. The P3 pellet contained little thioflavin S positive material and was not studied further. Isolated amyloid matrix from **C)** caput P2 and P4 and **D)** cauda P4 were stained with the amyloid dye, Thioflavin S (ThS) or nucleic acid stains, Sytox Green, Hoechst, and TOTO-3 and imaged using a Zeiss Axiovert 200M microscope equipped with epifluorescence. Caput P2 and P4 were also treated with 5% SDS while cauda P4 was treated with 70% formic acid for 15 minutes at room temperature to facilitate unwinding of the structure before staining. Insets, control, mock-stained samples in the absence of ThS or nucleic acid stains. Scale bar, 20 µm.

**Figure 2.**
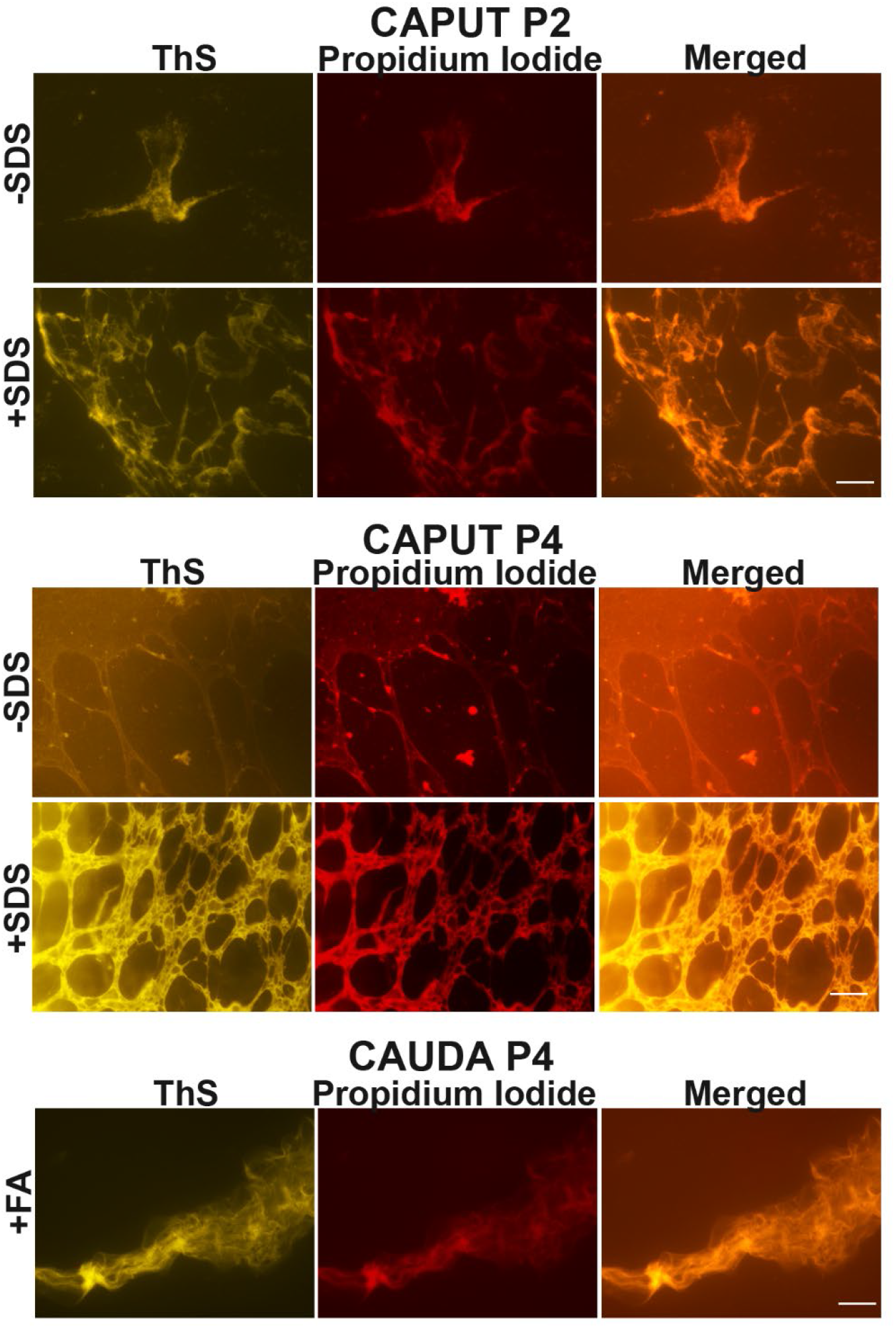
Extracellular nucleic acids colocalize with amyloid in the epididymal amyloid matrix. Caput P2 and caput and cauda P4 amyloids were stained with ThS to detect amyloid followed by propidium iodide to detect nucleic acid. Some caput P4 samples were treated with 5% SDS for 15 minutes while some cauda P4 samples were treated with 70% formic acid (FA) for 15 minutes prior to staining with ThS and propidium iodide. Scale bar, 20 µm.

To confirm the presence of nucleic acids and to determine the size associated with the P2 and P4 amyloids, we analyzed the samples by agarose gel electrophoresis and ethidium bromide (EtBr) staining. Caput P2 amyloid contained several populations of nucleic acids; that which was of high molecular mass and did not enter very far into the gel (red arrow), those between approximately 1.5 and 5 kb, and at 250 bp (Fig 3A). Similar nucleic acid populations were detected in the caput P4 amyloid but overall, there was less EtBr staining compared to caput P2. Because P4 may be a more highly ordered amyloid than P2 it may not completely enter the gel or unwind sufficiently to allow robust EtBr staining. Caput luminal fluid that did not undergo centrifugation (total) showed similar nucleic acid forms as in the caput P2 and P4 amyloids (Fig 3A). The supernatant fraction (S) obtained from the final centrifugation step to generate the P4 pellet did not exhibit any EtBr staining showing nucleic acids are only present in the particulate fraction. In the cauda epididymis the predominant populations of nucleic acid in the P2, P4 and total amyloid fractions were that which were of high molecular mass and a population at 250 bp (Fig 3A).

**Figure 3.**
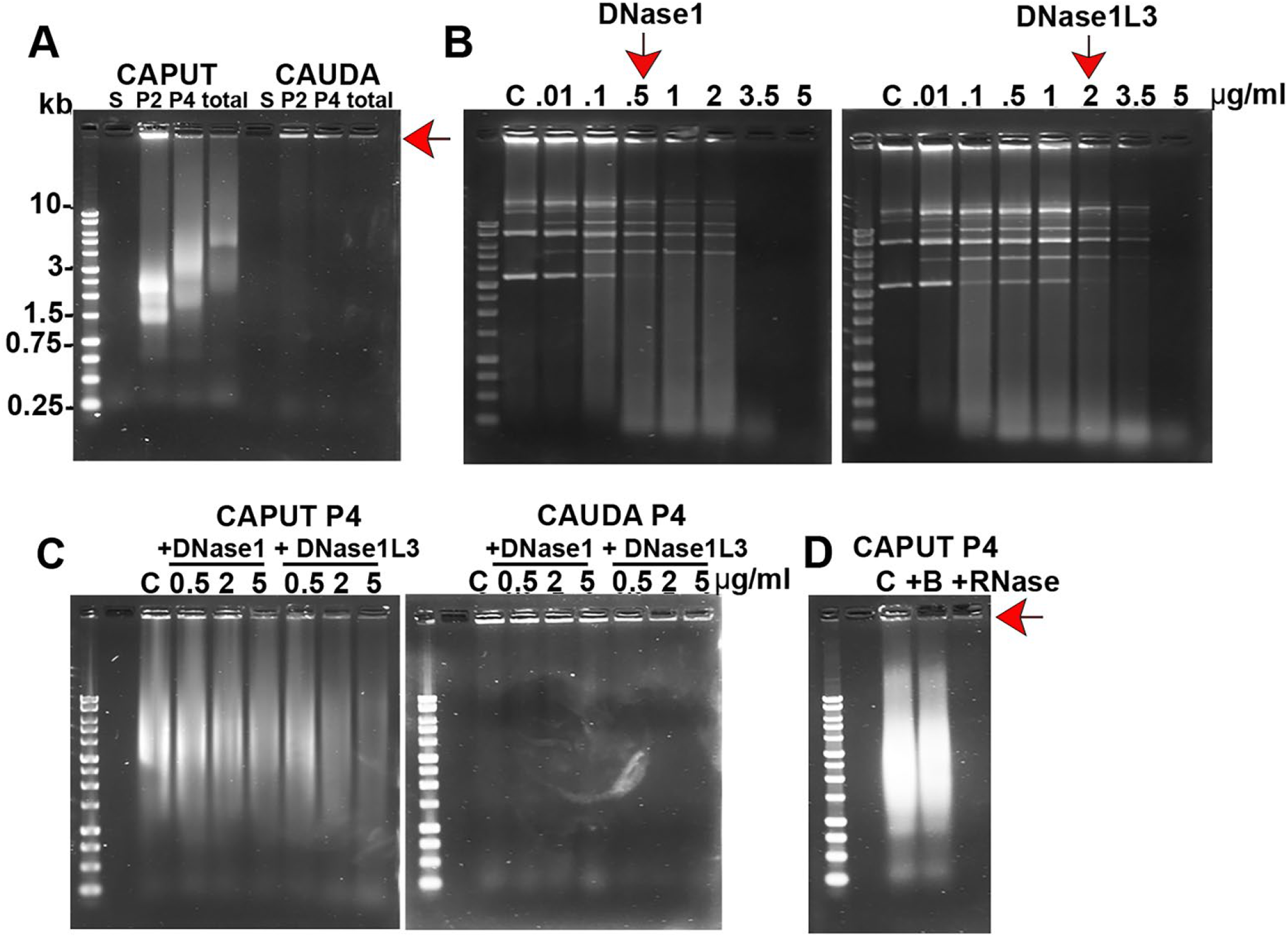
The epididymal amyloid matrix contains distinct populations of eDNA and eRNA. **A)** Twenty µgs of supernatant (S4/S), P2, P4 amyloid fractions and total luminal fluid (total) isolated from caput and cauda epididymis were separated on a 1% agarose-TAE gel and stained with EtBr to visualize nucleic acid. Red arrow indicates high molecular mass population of nucleic acid. **B)** 500 ng of plasmid DNA was incubated with 0.01-5 μg/ml DNase1 (left panel) and DNase1L3 (right panel) for 10 min at 37°C followed by 1% agarose gel electrophoresis and EtBr staining. Red vertical arrows indicate the DNase concentration at which ∼ 50% of the plasmid was digested. C, control, plasmid incubated in buffer only. **C)** Twenty µgs of caput P4 (left panel) and cauda P4 (right panel) amyloid fractions were incubated with 0.5, 2.5 and 5 µ/ml DNase1 or DNase1L3 and analyzed by agarose gel electrophoresis and EtBr staining. C, control, P4 amyloid incubated in the absence of DNase. **D)** Twenty µgs of caput P4 amyloid were incubated with DNase-free RNase for 30 min followed by agarose gel electrophoresis and EtBr staining. Red arrow indicates the high molecular mass population of DNA.

To determine if the EtBr staining in the epididymal amyloids represented eDNA, we incubated caput and cauda P4 amyloids with DNases and then determined the degree of degradation by agarose gel electrophoresis. DNase1 and DNase1L3 (DNase 1 like 3) are both Ca^2+^/Mg^2+^ - dependent endonucleases that participate in clearance of serum eDNA; however, DNase1L3 uniquely degrades antigenic forms of cell-free DNA, including DNA complexed with lipids and proteins(36). We used a plasmid degradation assay to compare the activity of DNase1L3 prepared in-house to that of a commercially purchased DNase1. Approximately 50% of plasmid DNA was digested by 0.5 µg/ml DNase1 while a similar percentage of plasmid digestion required 2 µg/ml of DNase1L3 (Fig 3B, arrows) suggesting the DNases have different activities or specificities. We then compared these two concentrations of DNase1 and DNase1L3 in their ability to digest the caput and cauda P4 amyloid matrix (Fig 3C). In addition, we compared the activity of both DNases at 5 µg/ml, a concentration at which the plasmid was completely digested by the two enzymes. Compared to control samples incubated in buffer, after 1 hour at 37°C DNase1L3 (2 µg/ml and 5 µg/ml) reduced the levels of the high molecular mass and 1.5-5 kb nucleic acid forms in the caput P4 amyloid but had no effect on the small 250 bp population (Fig 3C, left panel). Similarly, DNase1L3 digested some, but not all, of the high molecular mass nucleic acids in the cauda P4 amyloid (Fig 3C, right panel). In contrast, there was no digestion of nucleic acids in the cauda P4 amyloids after incubation with DNase1, and only a slight digestion of the caput P4 amyloid at its highest concentration of 5 µg/ml (Fig 3C, left panel). Together these results suggest eDNA is a component of the epididymal amyloid matrix and is present in several distinct populations including that which is resistant to, or protected from, DNase. The preferential ability of DNase1L3 to digest eDNA in the P4 amyloid over that of DNase1 is consistent with its known ability to degrade DNA complexed to protein (37).

We next determined if eRNA was present in the epididymal P4 amyloid. DNase-free RNase completely digested all the populations of nucleic acid in the caput epididymal P4 amyloid matrix except for the high molecular mass population (Fig 3D, red arrow). Together our results suggest the intermediate and small sized nucleic acids in the epididymal amyloids are RNA while the high molecular mass population is DNA. The slight digestion of the 1.5-5 kb population of nucleic acids by DNase1L3 suggests the enzyme may also possess some RNase activity in addition to its primary DNase functions (Fig 3C). Because the P4 amyloid preparations likely also contain extracellular vesicles/exosomes with DNA/RNA cargo we cannot rule out that some EtBr staining reflects these other populations of nucleic acids. It is also possible that some nucleic acids were released from the epididymal epithelial cells during the isolation of the amyloid fractions.

### Extracellular DNA and RNA are necessary for the maintenance of the epididymal amyloid matrix structure

Experiments were next carried out to determine if a function of eDNA is to maintain epididymal amyloid matrix structure. Caput P2 and caput and cauda P4 amyloids were incubated with 5 µg/ml DNase1L3 and stained with the nucleic acid stain Sytox Orange and ThS to determine if DNA digestion resulted in a change in the amyloid matrix structure. In this experiment cauda P4 was not pretreated with formic acid to unwind/expose the amyloid and therefore, compared to the caput, longer exposure times were needed during imaging of ThS and Sytox Orange to visualize the effects of DNase. For this reason, direct comparisons in ThS and Sytox Orange fluorescence intensities cannot be made between the different populations of amyloids from the different epididymal regions.

Caput P2 and caput and cauda P4 starting samples contained their characteristic sheets of ThS positive film (granular in P2) (Figs 4-6). Surprisingly, the addition of DNase reaction buffer alone affected the structure of the caput P2 and caput P4 amyloids. Specifically, the caput P2 amyloid starting sample, which typically exhibited a layer of small ThS positive granules, developed large holes/ clear zones suggesting points of perturbation and a local unwinding of structure (Fig 4A). An overall loss of amyloid was indicated by a significant decrease (p< 0.05) in total ThT levels, as determined by plate assay of the P2 sample exposed to DNase buffer compared to the starting sample (Fig 4B). Nucleic acids, as indicated by Sytox Orange fluorescence, also had a speckled pattern in the starting caput P2 sample (Fig 4A) and were reduced after exposure to buffer except around the periphery of the large holes where a thin edge of fluorescence was observed (SIFig 1). Because of this variable staining we were unable to measure a significant decrease in total nucleic acid levels in the P2 sample exposed to DNase reaction buffer compared to the starting sample (Fig 4C). Similarly, the caput P4 ThS positive amyloid film developed large holes, became fragmented, and exhibited reduced (p<0.05) amyloid levels after exposure to the DNase buffer (Fig 5A,B). Caput P4 also exhibited reduced nucleic acid levels when visualized by fluorescence microscopy except around the edges of the large holes where Sytox Orange fluorescence was bright (Fig 5A, SIFig1). DNase buffer had little effect on the cauda P4 amyloid; while the ThS stained amyloid appeared more web-like, these changes were typically only noticeable on the edges of the amyloid layer (Fig 6A) and therefore significant changes in total amyloid and nucleic acid levels were not observed (Fig 6B,C). Further, unlike the caput P2 and P4 amyloids, large holes with bright Sytox Orange fluorescent edges were rarely seen in the cauda P4 amyloid after exposure to DNase buffer.

**Figure 4.**
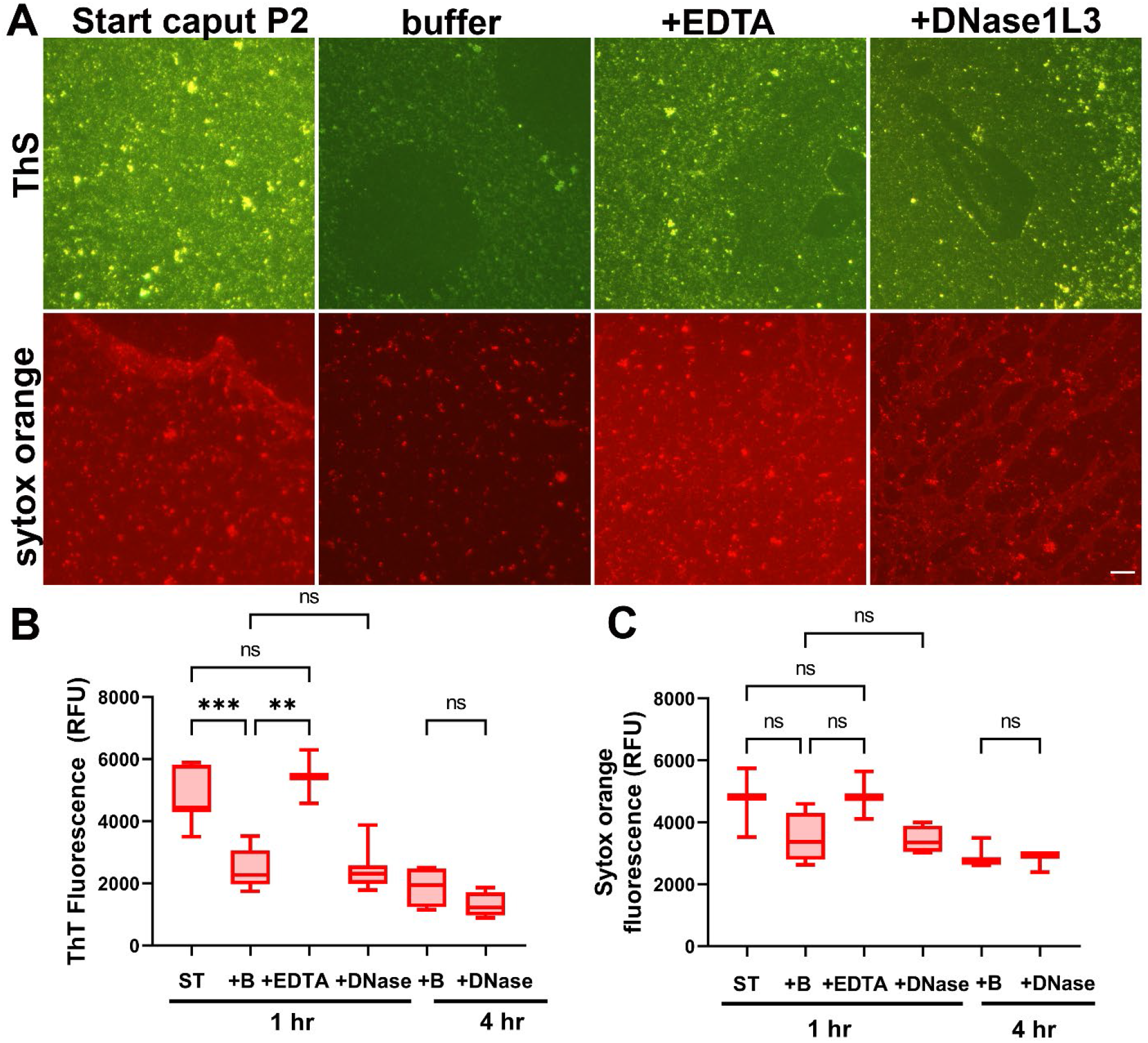
An endogenous nuclease activity partially disassembles the caput P2 epididymal amyloid matrix. **A)** Caput P2 amyloid fractions (30 μgs) were incubated with DNase buffer (buffer), DNase buffer + EDTA, or 5 µg/ml DNase1L3 for 1 hour at 37°C. Start sample was caput P2 amyloid incubated in buffer without MgCl_2_. Samples were dried overnight on a slide and incubated with ThS to visualize amyloid while a second set of samples were incubated with Sytox Orange to detect nucleic acids. Images were captured using a Zeiss Axiovert 200M microscope equipped with epifluorescence. Scale bar, 20 µm. A proportion of each sample analyzed in A) was added to a 96 well plate in the presence of either **B)** 20 µM ThT or **C)** 0.2 µM Sytox Orange and relative fluorescence units measured with a Tecan plate reader. Some samples were incubated for 4 hr in DNase1L3 or buffer only. n=3-8 experiments. Statistical analyses were performed using ANOVA followed by multiple comparisons test in GraphPad Prism 9.5. * p=0.01, **p=0.002.

**Figure 5.**
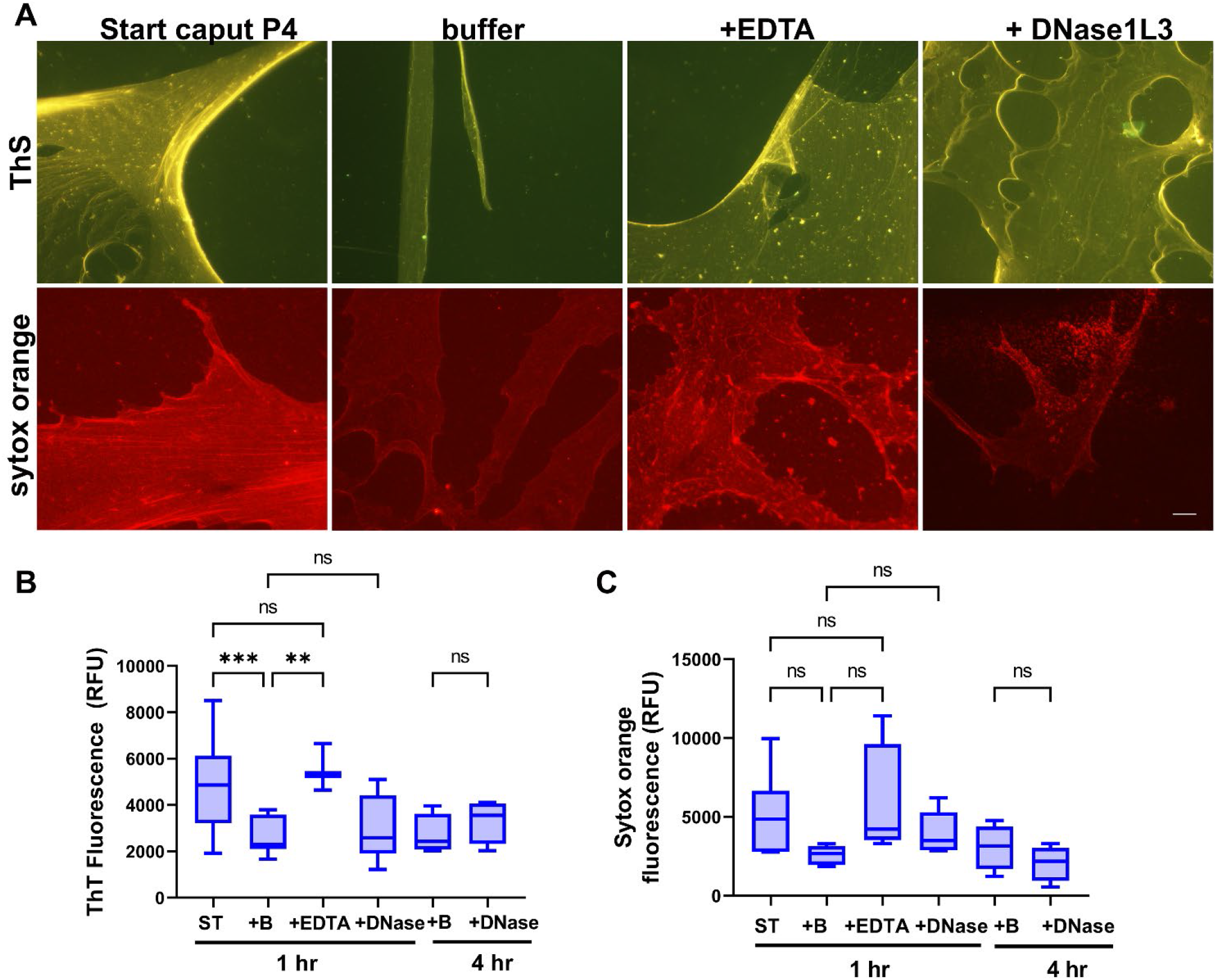
An endogenous nuclease activity partially disassembles the caput P4 epididymal amyloid matrix. **A)** Caput P4 amyloid fractions (30 ug) were incubated with DNase buffer (buffer), DNase buffer + EDTA, or 5 µg/ml DNase1IL3 for 1 hour at 37°C. Start sample was caput P4 amyloid incubated in buffer without MgCl_2_. Samples were dried overnight on a slide and incubated with ThS to visualize amyloid while a second set of samples were incubated with Sytox Orange to detect nucleic acids. Scale bar, 40 µM. A proportion of each sample analyzed in A) was added to a 96 well plate in the presence of either **B)** 20 µM ThT or **C)** 0.2 µM Sytox Orange and relative fluorescence units measured with a Tecan plate reader. Some samples were incubated for 4 hr in DNase1L3 or buffer only. n=3-9 experiments. Statistical analyses were performed using ANOVA followed by multiple comparisons test in GraphPad Prism 9.5. ** p=0.002, ***p= 0.0005.

**Figure 6.**
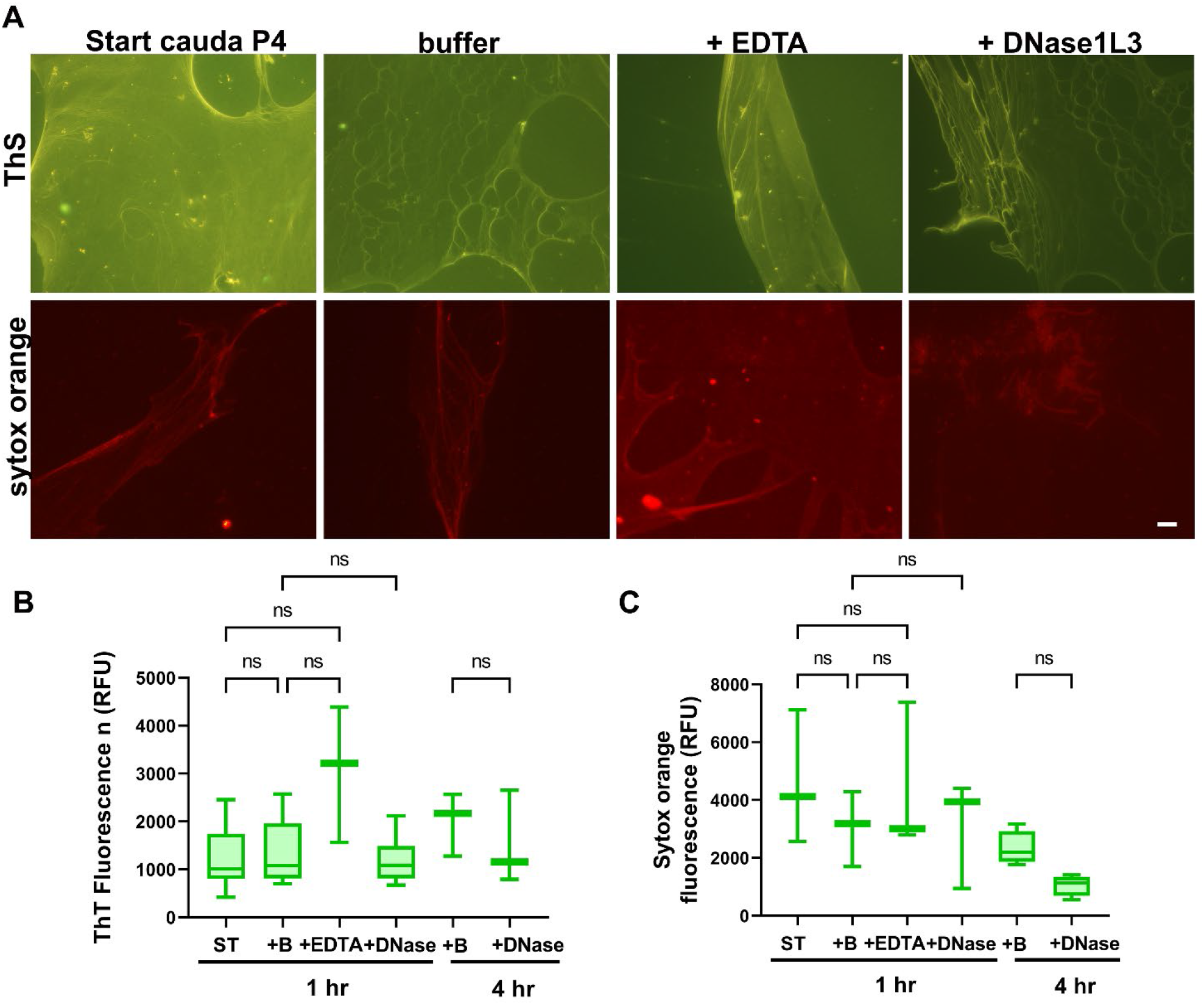
The cauda P4 amyloid matrix is resistant to disassembly. **A)** Cauda P4 amyloid fractions (30 ug) were incubated with DNase buffer (buffer), DNase buffer + EDTA, and 5 µg/ml DNase1L3 for 1 hour at 37°C. Start sample was cauda P4 amyloid incubated in buffer without MgCl_2_. Samples were dried overnight on a slide and incubated with ThS to visualize amyloid while a second set of samples were incubated with Sytox Orange to detect nucleic acids. Images were captured using a Zeiss Axiovert 200M microscope equipped with epifluorescence. Scale bar, 40 µM. A proportion of each sample analyzed in A) was added to a 96 well plate in the presence of either **B)** 20 µM ThT or **C)** 0.2 µM Sytox Orange and relative fluorescence units measured with a Tecan plate reader. Some samples were incubated for 4 hr in DNase1L3 or buffer only. n=3-9 experiments. Statistical analyses were performed using ANOVA followed by multiple comparisons test in GraphPad Prism 9.5., n.s., not significant.

The only difference between the starting P2 and P4 amyloid samples and those incubated with DNase buffer was the presence of 1 mM MgCl_2_. While we cannot rule out that it has a direct effect on the amyloid matrix, it is possible that MgCl_2_ activated an endogenous nuclease that had been previously detected in the mouse epididymal lumen and proposed to be DNase1L3 (38). Indeed, the addition of EDTA resulted in P2 and caput P4 amyloids that maintained their levels of amyloid and nucleic acids and exhibited structures comparable to that in the starting samples suggesting an inhibition of a MgCl_2_-dependent endogenous nuclease activity (Figs 4,5). Because of the effect of the buffer on the P2 and P4 amyloid and nucleic acid levels, we were unable to detect any additional changes in structure and in total amyloid and nucleic acid content after the addition of exogenous DNase1L3, even after a longer incubation time of 4 hr (Figs 4-6). The ring of Sytox Orange fluorescence around the buffer-induced holes remained, and even sometime appeared brighter (SIFig1). These results suggest that the exogenous DNase1L3 might help expose other populations of nucleic acids at sites where the amyloids have unwound/disassembled but whether these populations represent eRNA or DNase-resistant eDNA is not known.

Although the caput P2 and P4 amyloid films formed holes and fragments in the presence of DNase buffer (with and without DNase1L3) the amyloid matrix did not completely disassemble. These experiments further support that there may be several populations of eDNA associated with the amyloid matrix including those that are highly stable/resistant or inaccessible to DNase. Our results are consistent with those observed with the agarose gels showing high molecular mass DNA that remained following exposure to DNase1L3 (Fig 3). Although RNAsin was included during the incubations to prevent effects of possible confounding RNases on amyloid matrix dispersion, whether RNAsin would be a functional inhibitor of a potential RNase activity from DNase1L3 is not known. Another possible reason why the epididymal amyloid did not completely disassemble is that other structural components, in addition to eDNA, contribute to the integrity of the epididymal amyloid matrix.

Therefore, we next carried out experiments to determine if eRNA also has a functional role in maintaining the epididymal amyloid matrix structure. The addition of DNase-free RNase caused the caput P2 and P4 amyloid to become fragmented as indicated by patchy ThS staining, with notably reduced (p<0.05) amyloid and nucleic acid content (Fig 7A-C). No exogenous MgCl_2_ was included during the reactions and therefore samples exposed to buffer only served as controls since in these samples ThS, ThT, and Sytox Orange fluorescence were not different from that of starting P2 and P4 amyloids (data not shown). Although the effects of RNase on the structure of the cauda P4 amyloid were less pronounced than in the caput, some areas of the dense cauda P4 amyloid developed a swiss cheese appearance with small holes scattered throughout. However, while there was a loss of nucleic acids in the cauda P4 amyloid following RNase treatment, overall amyloid levels were not reduced (Fig 7A-C). Because RNase exposure resulted in significant changes in nucleic acid content in all epididymal amyloid samples (Fig 7) (p< 0.05) suggests that eRNA within the epididymal amyloid matrix may be more accessible to nucleases than eDNA.

**Figure 7.**
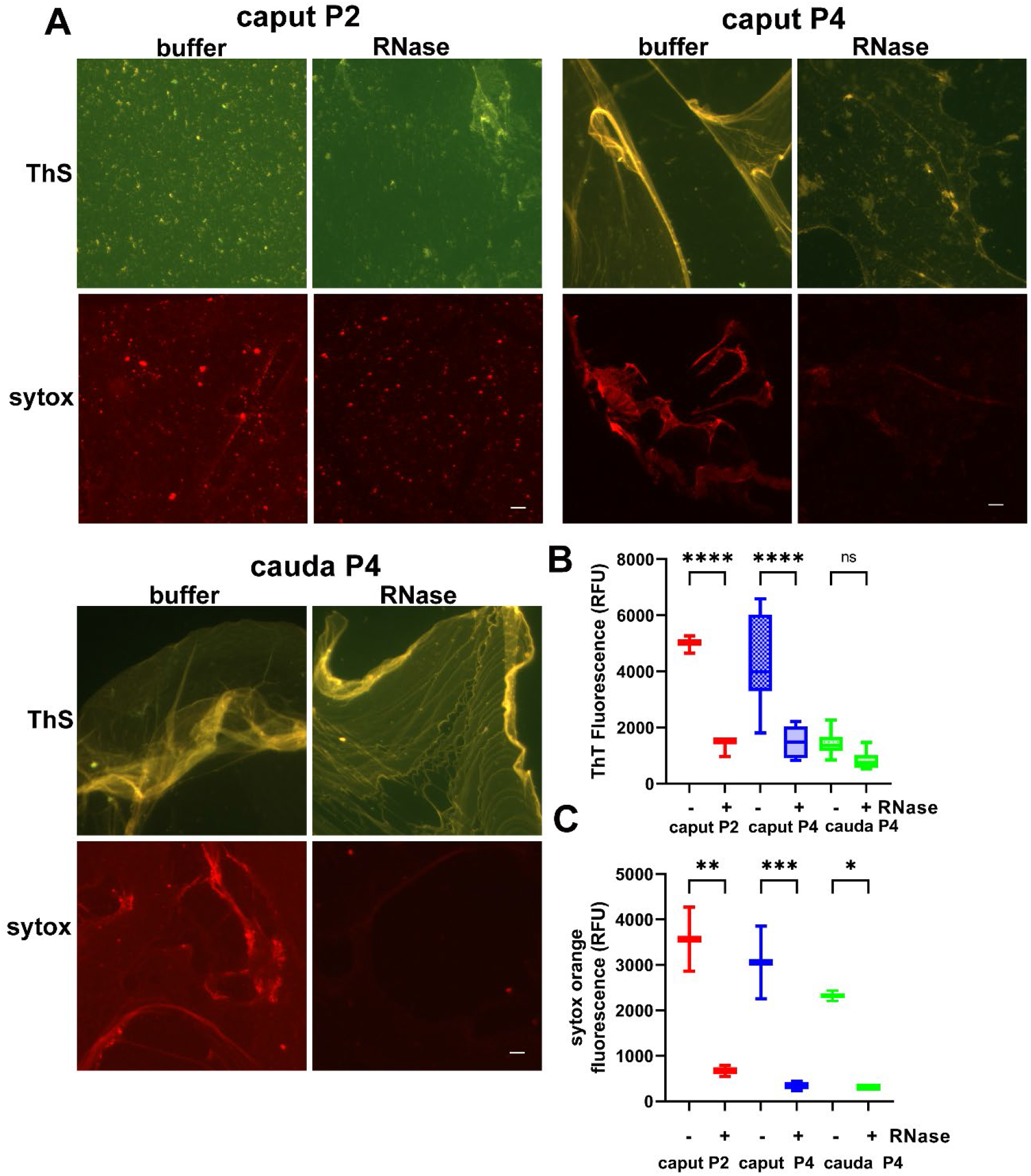
eRNA contributes to the maintenance of the epididymal amyloid matrix structure. **A)** Caput P2, caput 4 and cauda P4 amyloid fractions were incubated with DNase-free RNase or buffer only for 30 min at 37°C, dried overnight on a slide and stained with ThS or Sytox Orange to detect amyloid and nucleic acids, respectively. Scale bar, 40 µm. A proportion of each sample analyzed in A) was added to a 96 well plate in the presence of either **B)** 20 µM ThT or **C)** 0.2 µM Sytox Orange and relative fluorescence units measured with a Tecan plate reader. n= 3-4 experiments. Statistical analyses were performed using ANOVA followed by multiple comparisons test in GraphPad Prism 9.5. * p= 0.017, **p=0.002, *** p=0.0008, **** p<0.0001.

### DNase1L3 is present in the epididymal amyloid fractions

To determine if the endogenous nuclease activity detected in the caput and cauda epididymal amyloid fractions might be that of DNase1L3, we performed Western blot analysis using an anti-DNase1L3 antibody (a generous gift from P. Keyel, TTU). A 37 kDa protein characteristic of DNase1L3 monomer (Fig 8A), that was not detected with the control preimmune serum (Fig 8B), was present in all fractions from the caput epididymis including the soluble (supernatant) fraction. A minor ∼ 150 kDa immunoreactive band was also detected in the caput P4 amyloid. In the cauda epididymis the ∼150 kDa DNase1L3 isoform was present in equal, if not greater amounts, than the 37 kDa monomer; both of which were mostly in the P4 amyloid fraction. Together, these data suggest that DNase1L3 may form an SDS-stable tetramer via free cysteine residues or a complex with other proteins in the epididymal lumen. Indeed, based on its presence in the P2 and P4 fractions, DNase1L3 may be associated with, or be a part of, the epididymal amyloid matrix. If so, its activation could result in changes in the epididymal amyloid matrix structure presumably through its digestion of eDNA. Our observation that DNase buffer had little effect on the cauda P4 amyloid correlated with a change in the ratio of the DNase1L3 isoforms that were present (less monomer and increased higher molecular weight form) suggesting the two may be functionally related. However, further studies are needed to establish the role of the epididymal DNase1L3 in the epididymal amyloid matrix, including the significance of a possible 150 kDa isoform.

**Figure 8.**
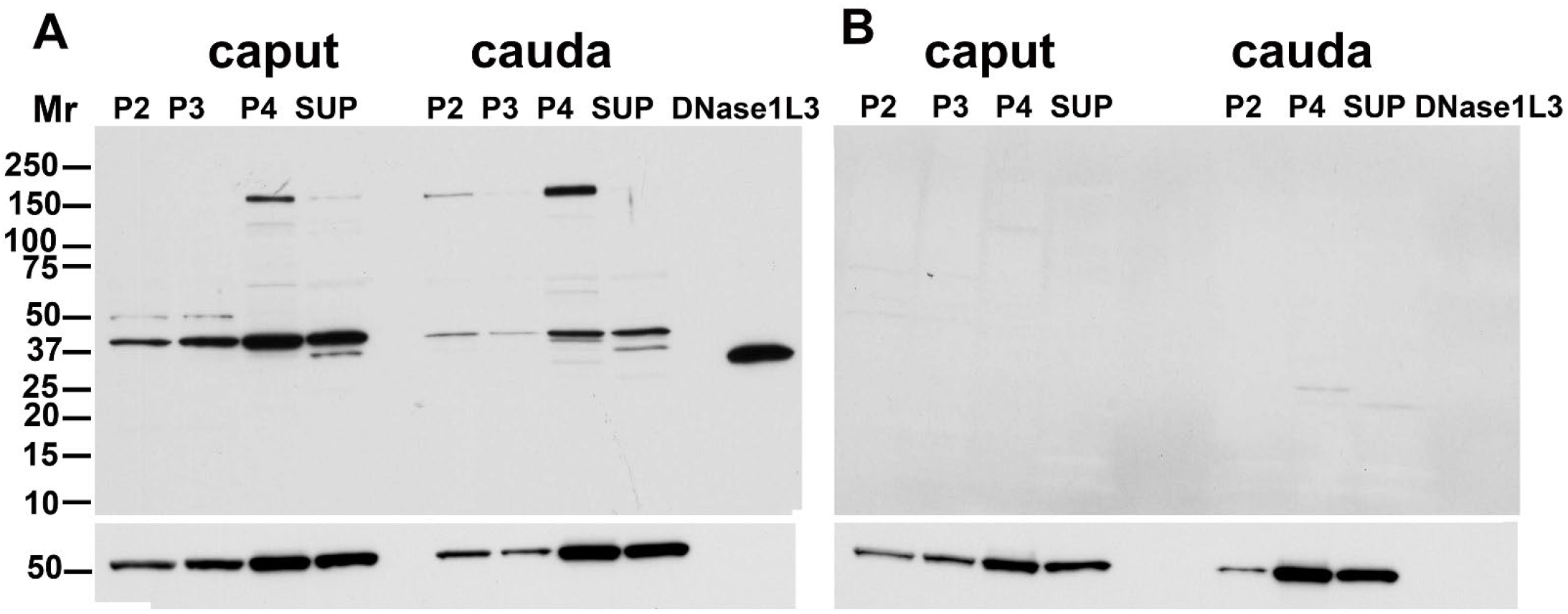
DNase1L3 is present in the epididymal amyloid matrix fractions. **A)** Western blot analysis of DNase1L3 in P2, P3, P4 and supernatant (S4,S) fractions from the caput and cauda epididymis. Human recombinant DNase1L3 was loaded as a positive control. **B)** Control Western blot analysis of the same samples in A) (except without cauda P3) using preimmune serum. Both blots were stripped and reprobed with α-tubulin as a loading control.

### Epithelial cells are a source of nucleic acids in the epididymal amyloid matrix

There could be several possible sources of the eDNA and eRNA in the epididymal amyloid matrix including release from the seminiferous tubule epithelium in the testis, epididymal epithelium, spermatozoa, and/or pathogens. Because the mice were healthy with no obvious sign of epididymal infection, the contribution from pathogens is likely minimal. We used the DC1 mouse caput epididymal cell line to determine if the epididymal epithelial cells produce an eDNA, eRNA-containing amyloid matrix. The use of a cell line allowed us to establish the presence of nucleic acids without the risk of contributions from spermatozoa. Cells were grown to confluency, conditioned media collected, and fractions containing P4 amyloids isolated as done with the epididymal amyloid matrix. As shown in Fig 9A, the P4 fraction stained with Hoechst and ThS, indicating the presence of nucleic acid associated with the amyloid. DC1 cell conditioned media, and a P4 fraction isolated from conditioned media, showed a high molecular mass population of nucleic acids that did not enter far into an EtBr-stained gel, faint levels of intermediate (1.5-5 kb) forms and a smaller (∼250 bp) form, populations like that observed in the P2 and P4 fractions from the caput epididymis (Fig 9B). Likewise, RNase caused the loss of the intermediate and smaller forms of nucleic acid with no effect on the high molecular mass population. Our results show the epididymal cell line mimics the epididymal epithelium and produces several distinct populations of nucleic acids that are components of an extracellular matrix: a high molecular mass population of eDNA and intermediate and small sized populations of eRNA. Thus, the epididymal epithelial cells are likely a source of eDNA and eRNA in the mouse epididymal amyloid matrix.

**Figure 9.**
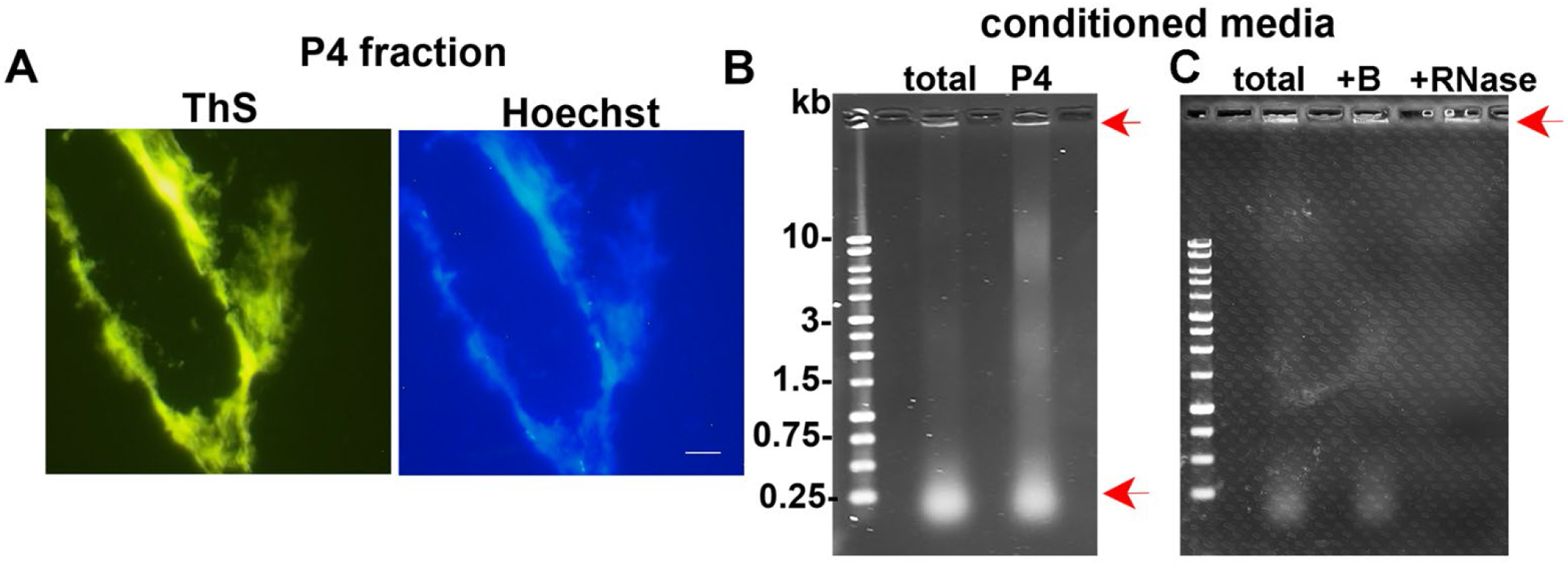
Epididymal epithelial cells are a source of nucleic acids in the epididymal amyloid matrix. **A)** The P4 amyloid fraction was isolated from the conditioned media from the DC1 caput epididymal epithelial cell line and stained with Thioflavin S to detect amyloid and Hoechst to detect nucleic acid. **B)** Twenty µgs of DC1 cell conditioned media (total) and the isolated P4 fraction were examined by 1% agarose-TAE gel electrophoresis and EtBr staining. Red arrows indicate the high molecular mass population of eDNA and the 250 bp population of RNA. **C)** Twenty µgs of total DC1 cell conditioned media were incubated with RNase or buffer only (B) for 30 min at 37°C and examined by agarose gel electrophoresis and EtBr staining. Red arrow indicates the high molecular mass population of eDNA.

### Polysaccharides are a component of the epididymal amyloid matrix and contribute to matrix integrity

To determine if the epididymal amyloid matrix contains mucin-like polysaccharides, caput and cauda P4 amyloids were stained with wisteria floribunda agglutinin (WFA) which is used to identify N-acetylgalactosamines beta 1 (GalNAc) residues in mucin-like glycoproteins, including chondroitin sulfate proteoglycans (39). Our experiments revealed that both the caput and cauda amyloid matrix contain complex GalNAc - containing sugars with the cauda P4 containing more GalNAc sugars compared to that in the caput P4 amyloid (Fig 10A, control). We next treated caput and cauda P4 amyloids with amylase, a glycoside hydrolase which breaks down the α-bonds of large α-linked polysaccharides into smaller oligosaccharides, and which is commonly used to digest bacterial biofilms (40). Amylase digestion resulted in a decrease in WFA staining and a fragmentation of the WFA-stained caput and cauda P4 amyloid matrix confirming the presence of polysaccharides and their contribution to maintaining overall structure (Fig 10A). More profound effects of amylase were observed in the caput P4 amyloid compared to the cauda P4 amyloid, consistent with it containing less polysaccharide and thus possibly being more easily dispersed. In addition, the caput P4 amyloid fraction exhibited a significant decrease (p<0.05) in amyloid content after exposure to amylase indicating a close association between the polysaccharide and amyloid components of the epididymal amyloid matrix (Fig 10B). A significant decrease in amyloid was not observed in the cauda P4 amyloid fraction following amylase exposure, possibly due to greater amounts of polysaccharide protecting the amyloid and/or its higher ordered state (Fig 10B).

**Figure 10.**
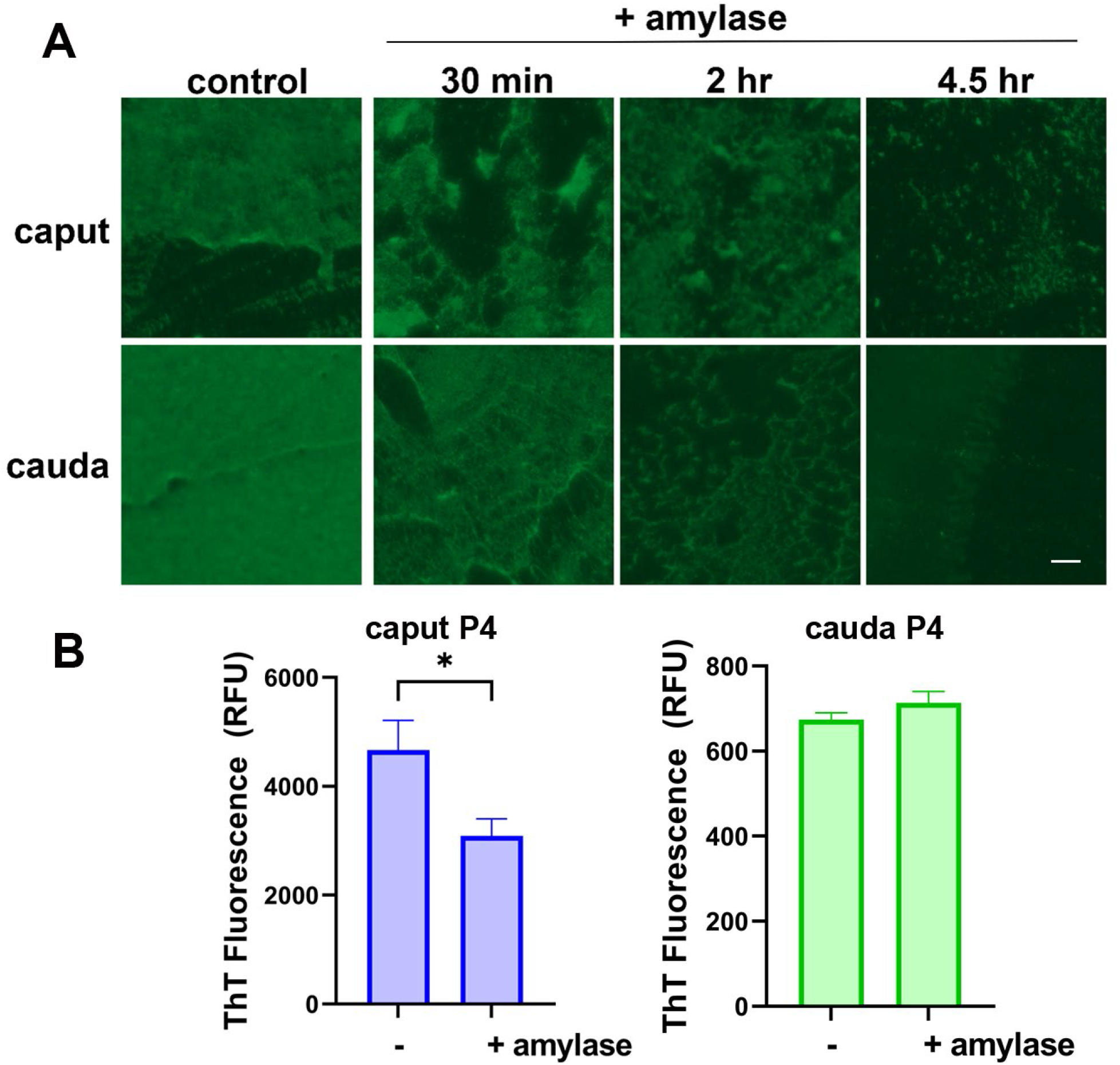
Mucin-like polysaccharides contribute to epididymal amyloid matrix integrity. **A)** Caput and cauda P4 amyloid fractions were dried on a slide overnight and stained with wisteria floribunda agglutinin (WFA), a stain for N-acetylgalactosamine beta 1 residues, before (control) and after treatment with 0.1% amylase. Scale bar, 20 μm. **B)** Caput and cauda P4 amyloid fractions were incubated with 0.1% amylase and total amyloid levels determined by ThT fluorescence. n=3. The relative ThT fluorescence units of control and amylase treated were compared by t-test. *, p<0.05.

We also stained whole epididymal tissue sections with Alcian blue pH 2.5 to detect acid mucins *in situ* including those with sulfated, carboxylic, hyaluronic, and chondroitin residues. Like that seen with WFA, more Alcian blue staining was observed in the cauda compared to the caput region indicative of more acidic mucin polysaccharides in the distal epididymis (Fig 11A). Some epididymal sections were also counterstained stained with neutral red which helped define structures in the epididymal lumen. The counterstained sections, which had a brown staining (red+blue), revealed a thin, branched matrix in the caput epididymal lumen that became a granular layer surrounding the spermatozoa and that continued into the cauda epididymal lumen (Fig 11A). Some sections also showed luminal patches of acid mucin polysaccharides with few spermatozoa suggesting the amyloid matrix is heterogeneous with distinct layers of sperm and polysaccharides distributed throughout the lumen (Fig 11A, *). Interestingly, in the cauda epididymal tubules we observed sperm organized into distinct mucin positive tunnels giving a swirled appearance (SIFig 2). Whether a function of the mucin polysaccharide/amyloid matrix is to stratify the sperm into organized structures for storage in the cauda epididymis requires further study.

**Figure 11.**
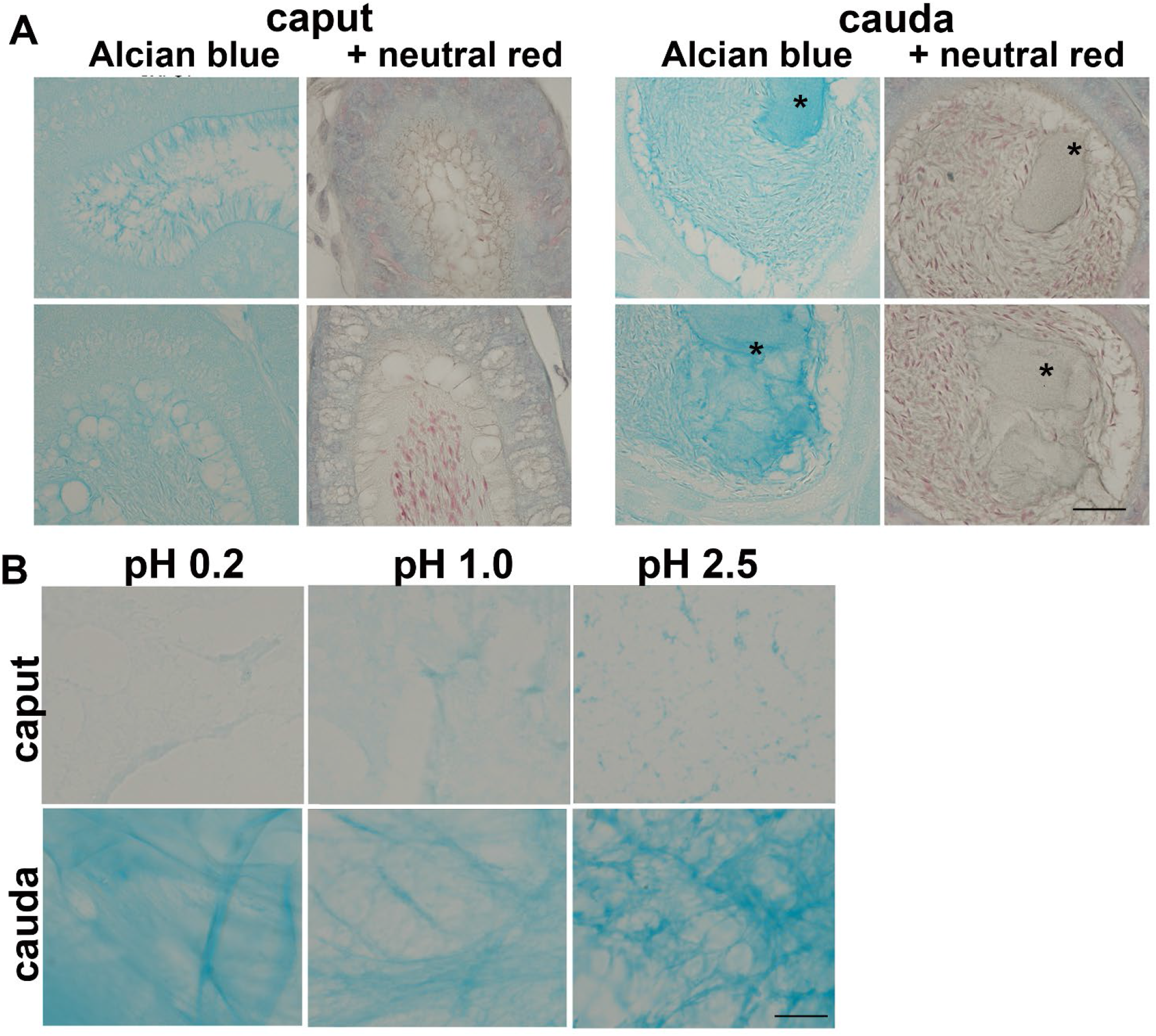
Acid mucin polysaccharides, including those strongly sulfated, are present in the epididymal amyloid matrix and enriched in the cauda epididymis. **A)** Mouse epididymal tissue sections were stained with Alcian blue, pH 2.5 with and without counterstaining with neutral red to detect acid mucins including those with sulfated, carboxylic, hyaluronic, and chondroitin residues. More robust staining was detected in the cauda compared to the caput epididymis. Neutral red stains nuclei and lysosomes and as a counterstain helped reveal the mucin-rich amyloid matrix (blue + red= brown) surrounding spermatozoa. * Area of polysaccharides without spermatozoa. **B)** P4 amyloid fractions isolated from the caput and cauda epididymis were stained with Alcian blue at pH pH 2.5 to detect total acid mucins, pH 1 to detect only sulfated acid mucins and pH 0.2 to detect strongly sulfated acid mucins. Scale bar, 20 μm.

P4 amyloid fractions isolated from the caput and cauda luminal fluid were also stained with Alcian blue pH 2.5 to detect total acid mucins, pH1 to detect only sulfated acid mucins and pH 0.2 to detect strongly sulfated mucins (). Alcian blue staining was observed at all pH conditions suggesting several types of acid mucins, including those that are strongly sulfated, are components of the epididymal amyloid matrix (Fig 11B). Together our studies show that mucin polysaccharides are an integral part of the amyloid matrix and contribute to the maintenance of its structure. Further, higher levels of polysaccharides are a characteristic of the cauda epididymal amyloids, which may be reflective of it being a more mature and established structure.

## Discussion

### The epididymal amyloid matrix has the features of a bacterial biofilm

Here we show that the mouse epididymal amyloid matrix contains eDNA, eRNA and complex polysaccharides, components that are critical for maintaining the extracellular epididymal amyloid matrix structure much like their functions in bacterial biofilms. Our previous work showing the epididymal amyloid matrix has host defense functions revealed a biological role similar to that of biofilms (7). Together, our studies suggest evolutionarily conserved mechanisms underlie the assembly and function of highly specialized extracellular matrices (ECMs) that protect a bacterial community and the mammalian germ line. Other mammalian host defense structures such as gut α-defensin nanonets form an amyloid/eDNA structure to enable trapping of the pathogen providing additional evidence of how nature has conserved some of the features of biofilm in higher organisms (41). However, whether eRNA and polysaccharides are also functional components of these structures is not known. Nanonets are also distinct in that they form solely in response to a pathogen while the epididymal amyloid matrix is always present, including in the absence of infection, implying roles in addition to host defense.

### eDNA and eRNA are integral components of the epididymal amyloid matrix and exhibit behaviors like their counterparts in bacterial biofilm

We show that one role of the eDNA in the epididymal amyloid is to support the overall integrity of the structure since upon activation of an endogenous nuclease activity (possibly DNase1L3) the amyloid matrix structure developed large holes and became fragmented. Disruption of the amyloid matrix structure was noted especially in the caput P2 and P4 samples consistent with it being an immature structure still undergoing assembly in this region. The caput amyloid structure was also affected by exogenous RNase indicating both eDNA and eRNA are part of the epididymal amyloid matrix and required for maintenance of its structure. Although the discovery of eRNA in bacterial biofilms is relatively recent (42,43), there is evidence of DNA/RNA-dependent biofilms. Studies in *Pseudomonas* in which eDNA was isolated from biofilms under conditions that maintained its *in vivo* biophysical properties revealed that purine-rich RNA was also present in the eDNA network (25). These studies suggested that non-canonical base pair interactions between the RNA and DNA allowed the formation of a nucleic acid gel and the viscoelastic properties of the biofilm.

eDNA likely performs additional roles in the epididymal amyloid matrix as it does in bacterial biofilms. eDNA has been shown to template the assembly of amyloidogenic proteins, suggesting it may serve as an initiator in the formation of an amyloid infrastructure(44). We have preliminary evidence that CRES, a component of the epididymal amyloid matrix, assembles into amyloid in the presence of DNA, a process that we speculate may initiate or contribute to the building of the structure (Cornwall unpublished). If so, some populations of eDNA may be integrated into the amyloid matrix such that they are partially protected from complete digestion or may be resistant to DNases. The cauda epididymal P4 amyloid, which our studies indicate is an advanced/mature amyloid structure compared to that in the caput epididymis, showed little to no changes in its structure in the presence of nucleases, a behavior comparable to the mature forms of many bacterial biofilms (19,45). The similarity in behaviors between the cauda P4 amyloid and mature biofilms suggests common mechanisms for their enhanced stability.

The eDNA in some biofilms has been shown to be like the intracellular DNA but with different methylation patterns (46). However, other studies indicate biofilm eDNA has a different higher ordered structure than chromosomal DNA and often is in a stabilized noncanonical configuration. In *E.coli* and other bacteria, eDNA was shown to have a Holliday junction-like arrangement that was critical for biofilm structural and mechanical integrity (20). Studies of *P. aeruginosa* eDNA showed that it formed G-quadruplexes, noncanonical secondary structures formed by G-rich sequences that contribute to the stability of the biofilm (47). In some bacterial biofilms eDNA changes its structure from the right-handed nuclease sensitive B-DNA form, the most common DNA conformation in biological conditions, to a more stable and nuclease resistant left-handed Z -DNA conformation as the biofilm matures (45). Altogether, the distinct and highly ordered structures of eDNA, and possibly also eRNA, in biofilms are thought to be mechanisms to allow the maturing biofilm to become more resistant to host responses providing further stability and protection for the bacterial colony. While studies are ongoing in our lab to determine if eDNA in the cauda P4 amyloid is in the Z conformation and/or has formed G-quadruplexes, it is possible such a transition might occur to enhance the stability of the amyloid matrix in the cauda epididymis which performs a critical function as a storage site for the mature spermatozoa.

The populations of eDNA and eRNA that we observed in the epididymal amyloid matrix exhibit sizes comparable with what has been detected in some biofilms; these include a high molecular mass population of eDNA (> 10 kb) that does not migrate well into the gel and an eRNA population of ∼ 250 bp (42). In other biofilms, however, smaller eDNA fragments have been observed. In a hyperbioflm-forming clinical variant of *P. aeruginosa* the release of eDNA fragments (100 bp-12000bp) from a subpopulation of the bacteria enabled the biofilm to become resistant to DNase I unlike the wildtype cells that released typical > 10 kb eDNA forms (48). These data suggest that eDNA itself may change in biofilms as an adaptive response to the environment resulting in an altered biofilm structure. Whether the >10 kb forms of eDNA observed in the epididymal amyloid matrix would change following an infection or an exposure to other environmental stressors, and if, and how, these changes might contribute to a changed epididymal amyloid matrix structure and function requires further investigation. In our previous studies, we observed that the epididymal amyloid matrix formed different amyloid structures with different host defense functions depending on the bacterial strain it encountered (7). These results indicate the epididymal amyloid, like biofilms, has an inherent ability to respond to environmental cues, which in part could be mediated by eDNA.

### Source of nucleic acids in the epididymal amyloid matrix

Although we found that epididymal epithelial cells are a source of eDNA and eRNA in the amyloid matrix produced *in vitro*, we cannot conclude that this is the only source of eDNA and eRNA in the mouse epididymal amyloid matrix. In addition to the principal cells, the primary epithelial cell in the epididymis, there are other cell populations that could contribute nucleic acids including apical cells, clear cells, and basal cells. In addition, eDNA and eRNA could come from damaged spermatozoa or secretions from the testis and efferent ducts. Further studies are also needed to determine the mechanisms by which nucleic acids become extracellular in the epididymal lumen. The release of DNA from bacteria is still under investigation but can include a suicidal programmed bacterial apoptosis or a fratricide-induced death or excretion from living cells (49,50). eDNA in extracellular traps released by neutrophils (NETs) comes from that bound to histones through cell lysis and, occasionally, from mitochondrial DNA (51,52). In addition to eDNA in the epididymal amyloid matrix, the caput and cauda epididymal P4 amyloid preparations likely contain extracellular vesicles/exosomes (EVs) and condensates which may contain DNA and RNA (53,54). If so, EVs could be one mechanism by which epididymal cells provide eDNA and eRNA to the amyloid matrix; however, we think it likely other mechanisms are involved as well considering nucleic acids are a part of the amyloid matrix infrastructure.

### Polysaccharides contribute to the maintenance of the epididymal amyloid matrix structure

Our results show that polysaccharides are a component of the amyloid matrix throughout the epididymis with higher amounts present in the mature epididymal amyloid structure. Using wisteria floribunda agglutinin and Alcian blue staining we showed in the cauda epididymis and in isolated cauda P4 amyloid fractions robust levels of N-acetylgalactosamines beta 1 (GalNAc beta 1-3 Gal) residues that are present in a variety of glycoconjugates including sulfated and acid mucin-like polysaccharides. Similarly, sulfated glycosaminoglycans, alginate and other negatively and positively charged extracellular polymeric substances are key constituents of biofilms from a variety of environments (27,55,56). These mucin-like sugars help create the stickiness or “glue” that holds the biofilm together. We observed the cauda epididymal P4 amyloid likewise formed a sticky pellet during isolation that may be due to its mucin-like polysaccharides. Our data also showed that amylase treatment disrupted the caput P4 epididymal amyloid matrix indicating a role for polysaccharides in maintaining its integrity, like polysaccharide function in biofilms. The cauda P4 amyloid, however, was less affected by the amylase treatment than the caput possibly due to its higher polysaccharide content preventing significant change in amyloid content in the timeframe of the amylase exposure. Alternatively, a more stable amyloid/eDNA/eRNA infrastructure in the cauda P4 epididymal amyloid prevented its disassembly.

Mucin-like sugars also contribute to the protective nature of biofilms by forming physical barriers and providing a tolerogenic environment for the immunogenic bacteria (57). The epididymis must also maintain a tolerogenic environment for the naturally autoantigenic sperm. Although the mechanism by which the epididymis maintains tolerance is not known, our studies here showing parallels between the epididymal amyloid matrix and biofilms suggest that the thick mucin-like polysaccharide layer in the epididymal amyloid matrix could be integral for this protection.

### DNases in the epididymis and seminal fluid: regulation of epididymal amyloid matrix assembly/disassembly?

DNases in seminal fluid have been used as a marker for fertility in large mammals, as increased levels of DNase are correlated with a higher rate of successful fertilization (58). However, the fact that nature would intentionally surround spermatozoa, whose sole function is to provide DNA for the next generation, with DNases seems counterproductive. Also surprising, was our data showing DNase1L3 is present in the epididymal lumen, possibly associated with the epididymal amyloid matrix. We speculate DNases including DNase1L3 and possibly others have several roles in the male reproductive tract. Because eDNA is known to template amyloid assembly, DNase1L3 may be present locally within the epididymal amyloid matrix to control the assembly process by regulating the amount or location of the template eDNA. DNase1L3 digestion could also be a means to produce smaller fragments for remodeling the epididymal amyloid matrix for functional purposes. DNase(s) in seminal plasma include types I and II (59) such as bovine sperm fertility associated antigen that has 88% identity with human DNase1L3 (60). Seminal plasma DNases may help disassemble the epididymal amyloid matrix after ejaculation allowing the sperm to swim up the female tract for fertilization. Indeed, cell-free DNA, single and double-stranded RNA, and small RNAs (miRNA and piRNA) have been found in human semen (61–64). Whether the nucleic acids represent that released from digested epididymal amyloid matrix, secreted from other male tract organs such as prostate or seminal vesicle for distinct biological roles, or a combination thereof remains to be determined. In addition to DNase activity, seminal fluid contains amylase and α-glucosidase, both of which have been used as fertility markers and are known to assist in the liquefaction of semen (65,66). Amylase treatment has also been used to increase fertilization rates in men (67). Although, α-glucosidase is made and stored in the epididymis it does not appear to be active until the seminal fluid, which would be appropriate if its role was to help disassemble the epididymal amyloid matrix (66). Thus, while in our studies DNase1L3, RNase, and amylase, individually only partially disrupted the caput epididymal amyloid matrix with less influence on the cauda amyloid matrix, their combined activities in the seminal plasma might be sufficient to disassemble the structure and allow the release of spermatozoa. If so, higher levels of DNase and/or amylase in semen would likely result in increased fertility because of enhanced amyloid matrix disassembly.

Together, our studies show structural and functional relationships between a specialized ECM that surrounds the maturing mammalian spermatozoa and an ECM that surrounds a bacterial community suggesting evolutionarily conserved mechanisms for their assembly and functions. In addition to providing clues of how the epididymal amyloid matrix protects the germ line from pathogens, knowledge of how biofilms interact with and “nurture” a bacterial community may be helpful for understanding basic mechanisms of sperm maturation.

## Materials and Methods

### Isolation of epididymal amyloid fractions

All animal studies were conducted in accordance with the NIH Guidelines for the Care and Use of Experimental Animals using protocol #94041 approved by the Texas Tech University Health Sciences Center Institutional Animal Care and Use Committee. Male CD1 mice (22-28 weeks of age) were obtained from Charles River (Wilmington, MA) and maintained under a constant 12 h light/12 h dark cycle with food and water ad libitum. Nestlets were provided for nesting. Two mice were used for each experiment. The luminal contents from the caput and corpus-cauda epididymis were isolated by puncturing the tissue in Dulbecco’s PBS (dPBS) or 25 mM HEPES, pH 7.4, using a 30G needle and allowing material to disperse for 15 min at room temperature. The samples were centrifuged at 500 x g for 5 min to remove spermatozoa (P1). The S1 supernatant was spun again to remove any remaining spermatozoa and then centrifuged at 5000 x g for 10 minutes to generate pellet 2 (P2). The resulting supernatant (S2) was centrifuged at 15000 x g for 10 minutes to generate pellet 3 (P3) followed by centrifugation of the S3 supernatant at 200000 x g for 70 min using a tabletop Beckman ultracentrifuge to generate the final S4 supernatant (SUP) and P4 pellet (P4). All pellets were resuspended in dPBS or 25 mM HEPES, pH 7.4 and proteins quantified by BCA assay (ThermoScientific, Rockford, IL).

### Nucleic Acid and Thioflavin S staining

Caput P2 and P4 and cauda P4 samples were diluted 1:4 in dPBS and 5 µl were spread on slides (∼1cm x 1cm square) and allowed to dry overnight. Some of the dried samples were treated with 5% SDS or 70% formic acid for 15 minutes at room temperature. Slides were then washed 3x in H_2_O, 2 minutes each wash, before staining for nucleic acids and amyloid. For nucleic acids staining, slides were rehydrated with 25mM HEPES (Sytox Green) or dPBS (Hoechst, TOTO-3) for 2 minutes followed by incubation in 1µM Sytox Green (Invitrogen, Waltham, MA) in 25mM HEPES, 1µg/ml Hoechst 33342 in dPBS, or 10µM TOTO-3 (Molecular Probes, Eugene, OR) in dPBS for 15 minutes at room temperature in the dark. Mock-treated control slides were incubated in 25mM HEPES or dPBS. Slides were washed 3x in 25mM HEPES or dPBS followed by 1x ddH_2_O, 2 minutes per wash. Slides were mounted with Vectamount AQ (Vector Laboratories, Newark, CA) followed by imaging on a Zeiss Axiovert 200M microscope equipped with epifluorescence.

For Thioflavin S staining, slides were rehydrated with ddH_2_O for 2 minutes before incubation in 0.1% Thioflavin S in ddH_2_0 (filtered) (Sigma, St. Louis, MO) in a Coplin jar for 2 hours at room temperature in the dark. Mock-treated control slides were incubated in ddH_2_O without Thioflavin S. The slides were washed 3x in ddH_2_O and 3x in 50% ethanol, 10 minutes per wash. Slides were washed 1x in ddH_2_O followed by mounting with Vectamount AQ (Vector Laboratories, Newark, CA). For some experiments the ThS stained samples were then stained with propidium idodide. After the water wash, slides were equilibrated in dPBS by 3x 2 min washes and incubated with 20 µM propidium iodide for 15 min at RT in the dark. Slides were washed 3X in dPBS,1x ddH_2_0, 2 min each and mounted with Vectamount AQ. Images were captured on a Zeiss Axiovert 200M microscope equipped with epifluorescence.

### DNase and RNase treatment for ThS and Sytox Orange staining

Human DNase1L3 was prepared as described previously, aliquoted and stored at - 80°C(36). A fresh aliquot was thawed on ice, diluted to 0.2 mg/ml with dPBS or 25 mM HEPES, pH 7.4, and used immediately. Thirty μgs of caput P2, caput P4 and cauda P4 amyloid fractions in 25 mM HEPES, pH 7.4 were incubated in a final volume of 40 µl containing 20 units SupeRase-In RNAse inhibitor (Invitrogen), and dPBS (start samples). Buffer treated samples (buffer) also included 1 mM MgCl_2_, +EDTA samples included 1 mM MgCl_2_ and 5 mM EDTA; and +DNase samples included 1 mM MgCl_2_ and 5 µg/ml DNase1L3. All samples were incubated at 37°C for 1-4 hours. For RNase exposure, 30 µgs of caput and cauda amyloid fractions were incubated in a final volume of 40 µl containing dPBS and 1 µl (12.5 μg/ml final) DNase-free RNase in RNase buffer (10 mM Tris, pH 7.4, 5 mM CaCl_2_, 50% glycerol) (Sigma) or RNase buffer alone at 37°C for 30 min. Thirty-six µl were transferred to a 96 well plate and Thioflavin T (Sigma) added to 25 µM final or Sytox Orange (Invitrogen) added to 0.2 µM final and fluorescence measured after 5 min incubation at room temperature using a Tecan plate reader. ThT and Sytox Orange blank controls (all assay components except protein) were subtracted from the amyloid containing samples. Samples were done in duplicate or quadruplicate. The remaining 4 µl of the 40 µl total sample were spread on a slide, dried overnight, and stained with ThS, as described above, or Sytox Orange as follows. Slides were washed 3x in 25mM HEPES, 2 minutes per wash, incubated in 1µM Sytox Orange (Invitrogen, Waltham, MA) for 15 minutes at room temperature in the dark, washed 3x in 25mM HEPES, 1x in ddH_2_O, 2 minutes per wash and then mounted with Vectamount AQ (Vector Laboratories, Newark, CA) followed by imaging on a Zeiss Axiovert 200M microscope equipped with epifluorescence.

### Amylase treatment

One percent amylase was prepared in dPBS and sterile filtered (0.22µm) before enzyme activation in a 37°C water bath for 30 minutes. Immediately after enzyme activation, caput and cauda P4 were diluted 1:4 with 1% amylase (0.1% final) or dPBS (control) and reactions were carried out for 30 minutes at 37°C. Five µls of each sample were spread on to slides (∼1cm x 1cm square) and allowed to dry overnight prior to staining with WFA. Some samples were transferred to a 96 well plate, ThT added to 20 µM final and ThT fluorescence measured using a Tecan plate reader.

### Wisteria Floribunda Agglutinin Staining

Slides were rehydrated with dPBS for 2 minutes before incubation in 1:500 wisteria floribunda agglutinin (Invitrogen, Waltham, MA) overnight at room temperature in the dark in a humidified chamber. The slides were washed 3x in dPBS and 1x in ddH_2_0, 2 minutes per wash. Slides were mounted with Vectamount AQ (Vector Laboratories, Newark, CA) followed by imaging on a Zeiss Axiovert 200M microscope equipped with epifluorescence.

### Alcian blue staining

CD-1 mouse paraffin embedded epididymal tissue sections were deparaffinized through xylene (2 × 5 min each), rehydrated in ethanol (2 × 100%, 1 × 95% 2 min each) and water and stained with Alcian blue pH 2.5 (1% Alcian blue in 3% aqueous acetic acid, filtered) for 30 min in a Coplin jar. Slides were then put under running tap water for 2 min, rinsed in distilled water, dehydrated through ethanol (1x 95%, 2 × 100% 2 min each), cleared with xylene (1X 3 min) and mounted using Permount. Some tissue sections were counterstained with neutral red after Alcian blue staining. After the distilled water rinse, slides were incubated in 1% neutral red (in 0.1% acetic acid, filtered) for 1 min and then processed through ethanol and xylene as described for Alcian blue. For staining of isolated caput and cauda epididymal P4 amyloids, samples were spread on a slide and allowed to dry overnight. Slides were incubated in 1% Alcian blue, pH 2.5, pH 1.0 or pH 0.2 for 30 min, washed under running tap water (pH 2.5) or drained followed by a quick dip in distilled water (pH 1.0 and pH 0.2), dehydration through ethanol, xylene and mounting with Permount. 1% Alcian blue pH 1 was prepared in 0.1 N HCl while pH 0.2 was prepared in 10% sulfuric acid.

### Cell culture

The DC1 epididymal epithelial cell line (obtained from M. C. Orgebin-Crist (68) was cultured in 10 cm dishes (Nuncleon, ThermoScientific)) at 37°C in 5% CO_2_ in a humidified incubator in IMDM with phenol red supplemented with 10% fetal bovine serum (FBS) (R&D systems, Minneapolis, MN), 1mM Sodium Pyruvate, 0.1mM MEM non-essential amino acids, 4mM L-glutamine, 42 U/ml penicillin, 42 µg/ml streptomycin, and 0.001µM 5α-dihydrotestosterone (in 100% EtOH). To collect conditioned media, cells were grown until ∼70-80% confluency. Cells were washed 2x with warmed HBSS, 2 minutes per wash. Cells were then incubated in IMDM without phenol red and FBS but with all other additives at 37°C and 5% CO_2_ for 16 hours. Conditioned media was collected and spun at 500 × g for 5 minutes to remove cells. The conditioned media was moved to new tubes and spun again to remove any remaining cells. Conditioned media was then concentrated in a 3 kDa Amicon-Ultra-15 filter to a volume of 500 µl. Protein concentrations were determined by BCA assay (ThermoScientific, Rockford, IL). Different amyloid fractions were then prepared as done with luminal fluid from the epididymis. One hundred and seventy-five µg of concentrated conditioned media was centrifuged at 5000 × g for 10 minutes to generate pellet 2 (P2). The resulting supernatant (S2) was centrifuged at 15000 × g for 10 minutes to generate pellet 3 (P3) followed by centrifugation of the S3 supernatant at 200000 × g for 70 min using a tabletop Beckman ultracentrifuge to generate the final S4 supernatant (SUP) and P4 pellet (P4). Five µls of the sample were spread on slides and allowed to dry overnight before staining.

### Agarose Gel Electrophoresis

Twenty µg of supernatant (S4 supernatant, Fig 1B), P2, P4 and total (S1 supernatant, Fig 1B) from the caput and cauda epididymis and conditioned media from DC1 cells were diluted in dPBS to a final volume of 18 µl. Three µl of 6X loading buffer (Promega, Madison, WI) were added to each reaction and samples were run on a 1% agarose-TAE gel. Gels were stained with 2 µg/ml ethidium bromide (EtBr) for 20 minutes, washed in ddH_2_O for 30 minutes and imaged on an Azure Biosystem C300.

### Plasmid Digestion Assay

Five hundred ng of plasmid (pGEX-cs) were incubated with 0-5 μg/ml final DNase1 or DNase1L3 in 25 mM HEPES, pH 7.4, 0.5 mM MgCL_2_ in a reaction volume of 18 µl for 10 minutes at 37°C. Samples were run on a 1% agarose-TAE gel followed by staining with 2 µg/ml EtBr for 20 minutes and washed in ddH_2_O for 30 minutes. Gels were imaged on an Azure Biosystem C300.

### Epididymal Amyloid Matrix Degradation Assay

Twenty µgs of caput and cauda P4 were incubated with 3.6 µl of DNase1 or DNase1L3 (0.5, 2, or 5 µg/ml final concentration) in DNase storage buffer (20 mM HEPES, pH 7.4, 1mM CaCl_2_, 400 mM NaCl) and 4.5µl of 4X enzyme buffer (100 mM HEPES, pH 7.4, 2 mM MgCl_2_). dPBS was used to bring the reactions to a final volume of 18 µl. Control reaction contained 3.6 µl of DNase storage buffer in place of DNase and 4.5µl of 4X enzyme buffer. Reactions were incubated for 1 hour at 37°. Reactions were run on a 1% agarose-TAE gel, stained with 2 µg/ml EtBr for 20 minutes and washed in ddH_2_O for 30 minutes. Gels were imaged on an Azure Biosystem C300.

### Western blot analysis

Twenty-five μgs of caput and cauda P2, P3, P4 and final supernatant were separated on a 15% Criterion Tris-glycine gel (BioRad) followed by transfer to PVDF membrane as previously described (4). Blots were incubated in blocking buffer (3% milk/TBST (0.2% Tween-20) for 1 hr RT and incubated overnight at 4°C in blocking buffer containing 1:5000 rabbit anti-mouse DNase1L3 antiserum or preimmune serum. The blots were washed in TBST 3X followed by incubation in blocking buffer containing a goat anti-rabbit HRP tagged secondary antibody (Vector Labs) for 2 hr RT. Blot were washed repeatedly in TBST, incubated in Supersignal West Pico PLUS chemiluminescent substrate (Thermo Scientific) for 5 min and exposed to film. Blots were stripped using Restore PLUS Western blot stripping buffer (Thermo Scientific) and incubated as described above with a 0.5 μg/ml mouse α-tubulin antibody (clone B-5 1-2, Sigma) as a loading control.

## Notes

### Competing Interest Statement

The authors have declared no competing interest.

