## Supplemental Figures 1 and 2 for "The Mouse Epididymal Amyloid Matrix: A Mammalian Counterpart of a Bacterial Biofilm"

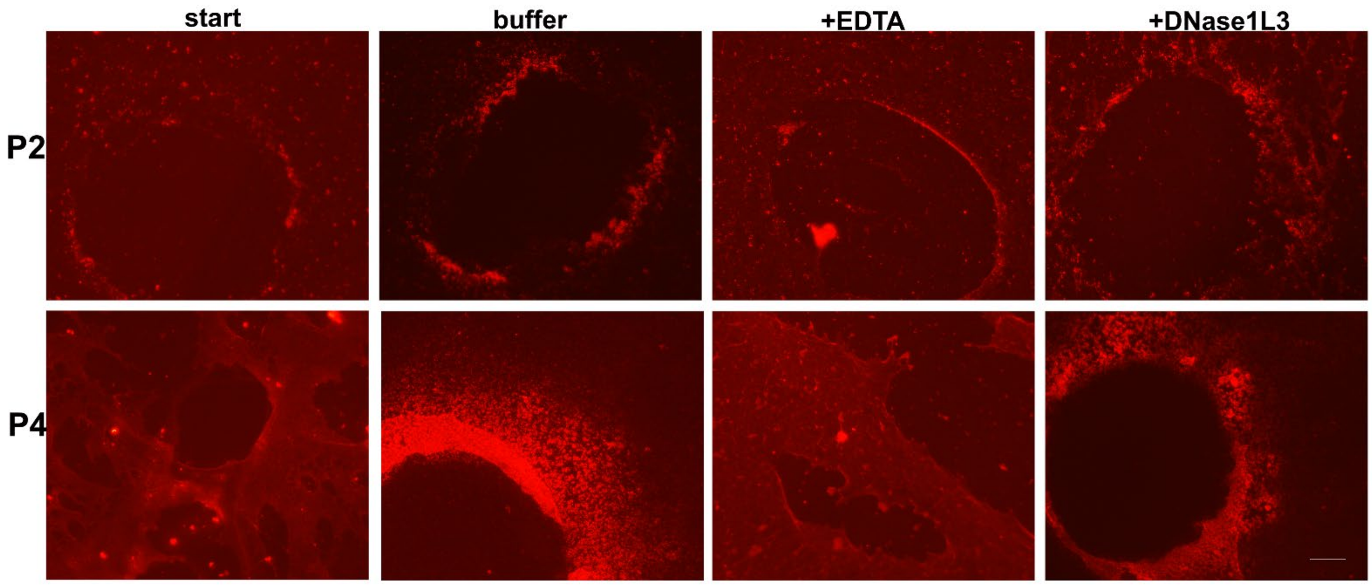


**SI Figure 1. Exposure of nucleic acids in the caput P2 and caput P4 amyloid matrix.** Images were captured from samples shown in Figs 4 and 5. Scale bar, 20 μm.

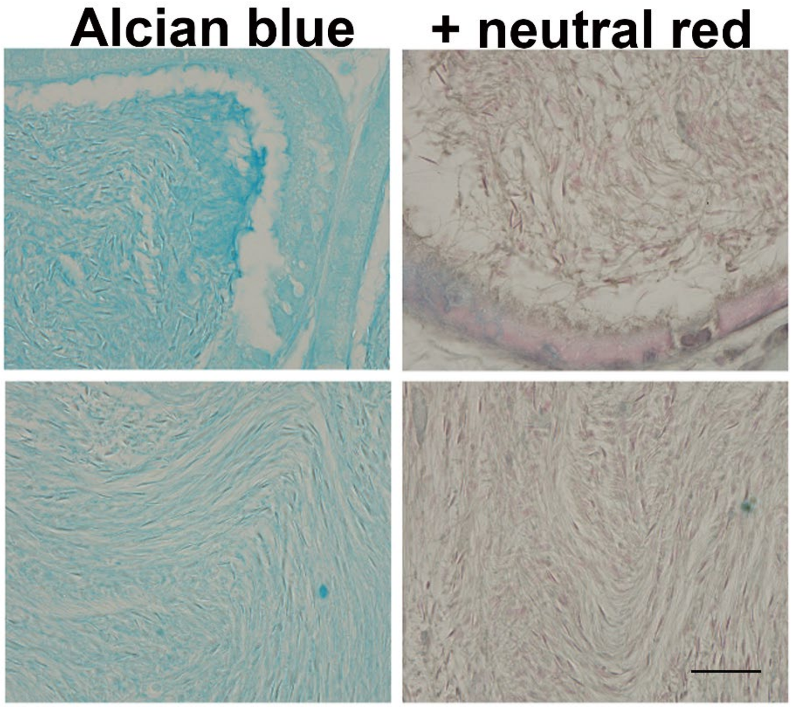


**SI Figure 2. Alcian blue and neutral red staining reveal stratified layers of spermatozoa in the cauda epididymis.** Images were captured from samples shown in Fig 11. Scale bar, 20 μm.
